# Putative mobilized colistin resistance genes in the human gut microbiome

**DOI:** 10.1101/2020.12.31.424960

**Authors:** Bruno G. N. Andrade, Tobias Goris, Haithem Afli, Felipe H. Coutinho, Alberto M.R. Davila, Rafael R. C. Cuadrat

## Abstract

**Background:** The high incidence of bacterial genes that confer resistance to last-resort antibiotics, such as colistin caused by MCR genes, poses an unprecedented threat to our civilization’s health. To understand the spread, evolution, and distribution of such genes among human populations, with the final goal of diminishing their occurrence in human environments should be a priority. To tackle this problem, we investigated the distribution and prevalence of potential mcr genes in the human gut microbiome we used a set of bioinformatics tools to screen the Unified Human Gastrointestinal Genome (UHGG) collection for the presence, synteny and phylogeny of putative mcr genes, and co-located antibiotic resistance genes.

**Results:** A total of 2,079 ARGs were classified as different MCR in 2,046 Metagenome assembled genomes (MAGs), present in 1,596 individuals from 41 countries, of which 215 MCRs were identified in plasmidial contigs. The genera that presented the largest number of MCR-like genes were *Suterella* and *Parasuterella*, prevalent human gut bacteria of which *Suterella wadsworthensis* is associated with autism. Other potential pathogens carrying MCR genes belonged to the genus *Vibrio*, *Escherichia* and *Campylobacter*. Finally, we identified a total of 22,746 ARGs belonging to 21 different classes in the same 2,046 MAGs, suggesting multi-resistance potential in the corresponding bacterial strains, increasing the concern of ARGs impact in the clinical settings.

**Conclusion:** This study uncovers the diversity of MCR-like genes in the human gut microbiome. We showed the cosmopolitan distribution of these genes in individuals worldwide and the co-presence of other antibiotic resistance genes, including Extended-spectrum beta-lactamases (ESBL). Also, we described mcr-like genes fused to a PAP2-like domain in *S. wadsworthensis*. Although these novel sequences increase our knowledge about the diversity and evolution of mcr-like genes, their activity and a potential colistin resistance in the corresponding strains has to be experimentally validated.

## Introduction

The prevalence of antibiotic resistance (AR) in clinical pathogens is a significant public health concern, especially in low and middle-income countries (LMICs)^1^. The misuse of antibiotics is the main driving factor for the rise of antibiotic-resistant bacteria. Still, its importance is often underestimated in community infections, as hospitalized infections gain the most attention^2^. Many previous studies have addressed the prevalence of AR in clinical environments^3 4 5^ and, recently, to also understand AR prevalence in non-hospitalized populations^6 7^. Most of these studies were conducted on cultivable clinical strains using microbiological methods that involve cultivation and antibiogram tests. However, the advances in high-throughput sequencing and bioinformatics enabled the study of metagenomes and access to the so-called human resistome.

The resistome is defined as the collection of the antibiotic resistance genes (ARGs) in a single microorganism, or in a microbial community, and has been investigated in different environments, such as soils^8^ or ocean^9^, and in host-associated microbiotas such as the animal^10^ or human^11^ gut. Understanding the human resistome in hospitalized and non-hospitalized populations are essential not only because the commensal microbiota can host and transfer ARGs from and to pathogenic bacteria by horizontal gene transfer (HGT)^12^, e.g., during an infection. In addition, HGT can also play a role in ARG mobilization to environmental communities by water and soil contamination^13^ or the food we ingest^14 15^. The gut microbiome is of particular interest in the investigation of ARGs in the human microbiota since it is the largest and most diverse^16^ and highly exposed to and affected by the intake of antibiotics. A potential influx of ARGs can occur via food intake and/or unhygienic conditions, and the efflux of ARGs to wastewater plants enhances the spread to other environments. As such, the human gut microbiome is thought to be responsible for transferring ARGs^17^ to the environment to a large extent. Therefore, the search for ARGs in the human gut microbiome, which is mostly performed using metagenomics and culturomics approaches, is one of the key fields to unravel the transfer of ARGs and the evolution of antibiotic resistance in bacteria. While past metagenomic studies on ARGs relied on shorter contigs, often below 50 kilobases, new assembly methods that allow the recovery of nearly complete bacterial genomes have been developed. Such methods have been applied to many studies regarding the human gut microbiome and allowed the recovery of thousands of metagenome-assembled genomes (MAGs) ^18–20^. These datasets were recently combined into one resource, the Human Gastrointestinal Bacteria Genome Collection (HGG)^21^, which makes this human gut MAGs collection a valuable resource to screen for ARGs. The advantage of MAGs versus traditional metagenome gene catalogs is manifold; the most apparent is the high accuracy of phylogenetic affiliations and often complete gene clusters, revealing gene synteny. Especially the latter is of high interest when studying ARGs since the genetic environment often shows the genetic mobility of ARGs, e.g., their location on genetic islands or plasmids^22^. Besides, it is also possible to investigate the presence of multi-drug-resistant (MDR) bacteria by detecting more than one ARG in the same bacterial genome or contig^9^ when using the MAGs approach.

Recently, colistin (Polymyxin E) has gained attention as the last line of defense drug against MDR bacteria, especially carbapenem-resistant gram-negative pathogens^23^. However, reports of colistin-resistant bacteria are becoming more frequent^24^, with its prevalence reaching as high as approximately 20%–40% among Carbapenem-Resistant *Klebsiella pneumoniae* (CRKP) in Italy and Greece^25^. In the past, the only known acquired resistance mechanism for colistin was mediated by chromosomal mutations, mainly in genes regulating the chemical additions of L-Ara4N and pEtN^26^. The first plasmid-mediated polymyxin resistance gene, designated mobilized-colistin resistance-1 (mcr-1), was described for *Enterobacteriaceae* in 2006^27^. Later it was followed by the additional mcrs, mcr-2^28^, mcr-3^29^, mcr-4^30^, mcr-5^31^ and, very recently, mcr-6 to mcr-10 ^32–36^. An intrinsic mcr-1-like homolog from *Moraxella osloensis* was described, named icr-Mo^3^7, raising the discussion about possible origins of MCRs in *Moraxella*. The spread of mcrs is of public health concern as it has been attributed to colistin's over-use, especially in livestock^38^ and aquaculture^39,40^. To unravel the presence of mcr-like genes we here screen the HGG for these genes and describe co-occurrence with other ARGs.

## Methods

### Data retrieval

We retrieved ~171 million non-redundant protein sequences (clustered at 100%) from the Unified Human Gastrointestinal Protein (UHGP) catalog^21^ plus the according to the metadata of the 286,997 corresponding metagenomic assembled genomes (MAGs). Also, we obtained the redundant protein identifiers mapped to the non-redundant protein representative for quantification. For the MAGs, including ARGs of interest, we retrieved the fasta sequences and GFF annotation files.

### Antibiotic resistance gene screening

To search for antibiotic-resistant genes in the human microbiome, we followed the methodology used in our previous study with oceanic samples 8. In short, we used the deepARG tool^41^ (model version 2), an in-depth learning approach developed to identify ARGs at both reads or operational reading frames level, to search for ARGs in the non-redundant proteins provided by the UHGP catalog. We then selected all proteins classified as mobilized colistin resistance (MCR) in the deepARG results and explored the prevalence of these putative ARGs in different countries and across diverse taxa.

### Plasmid classification

To verify if the putative genes are of chromosomal or plasmid origin, we applied the PlasFlow software^42^ with the default threshold of 0.7. This software uses neural network models trained on full genomes and plasmid sequences to predict the sequence origin with 96% accuracy.

### Phylogeny

Protein sequences classified as MCR present in contigs identified as plasmids were clusterized at 97% of sequence similarity with the software cd-hit v4.7^43^ to reduce the number of protein sequences in the tree. The representative sequences of each cluster (and reference sequences obtained at NCBI) were then submitted to NGPhylogeny.fr^44^ where the protein sequences were first aligned by Mafft^45^. The informative phylogenetic regions were selected by BMGE^46^, and the Maximum likelihood (ML) reconstruction was calculated by PhyML 3.0^47^ with the model selection performed by SMS (AIC method)^48^, and 100 bootstrap replicates to infer significance.

### Data visualization

For data visualization, we used jupyter notebook and python 3, with libraries pandas and matplotlib. For conserved domain visualization on unusual mcr-like sequences, we used NCBI blast and CDD (https://blast.ncbi.nlm.nih.gov/Blast.cgi), running Blastp with default parameters.

## Results and Discussion

We identified a total of 2,079 ARGs classified as MCR (13 MCR-1, 1 MCR-1.2, 9 MCR-2, 634 MCR-3, 456 MCR-4, 966 MCR-5) in 2,046 genomes (166 from isolates and 1880 from MAGs), present in 1,596 individuals (7.2% from the total 21,866 in the study) from 41 different countries. It is important to note the restricted classification of MCR in the deepARG model 2, only assigning up to MCR-5. The highest relative number of mcr-like genes (normalized by the total number of samples) was found in Haiti (>10%), followed by South Korea, Norway, and India (Figure 1). However, it is important to observe that a quantitative comparison is not possible due to the very unbalanced number of genomes from each country, varying from less than 50 to more than 50,000.

**Figure 1:**
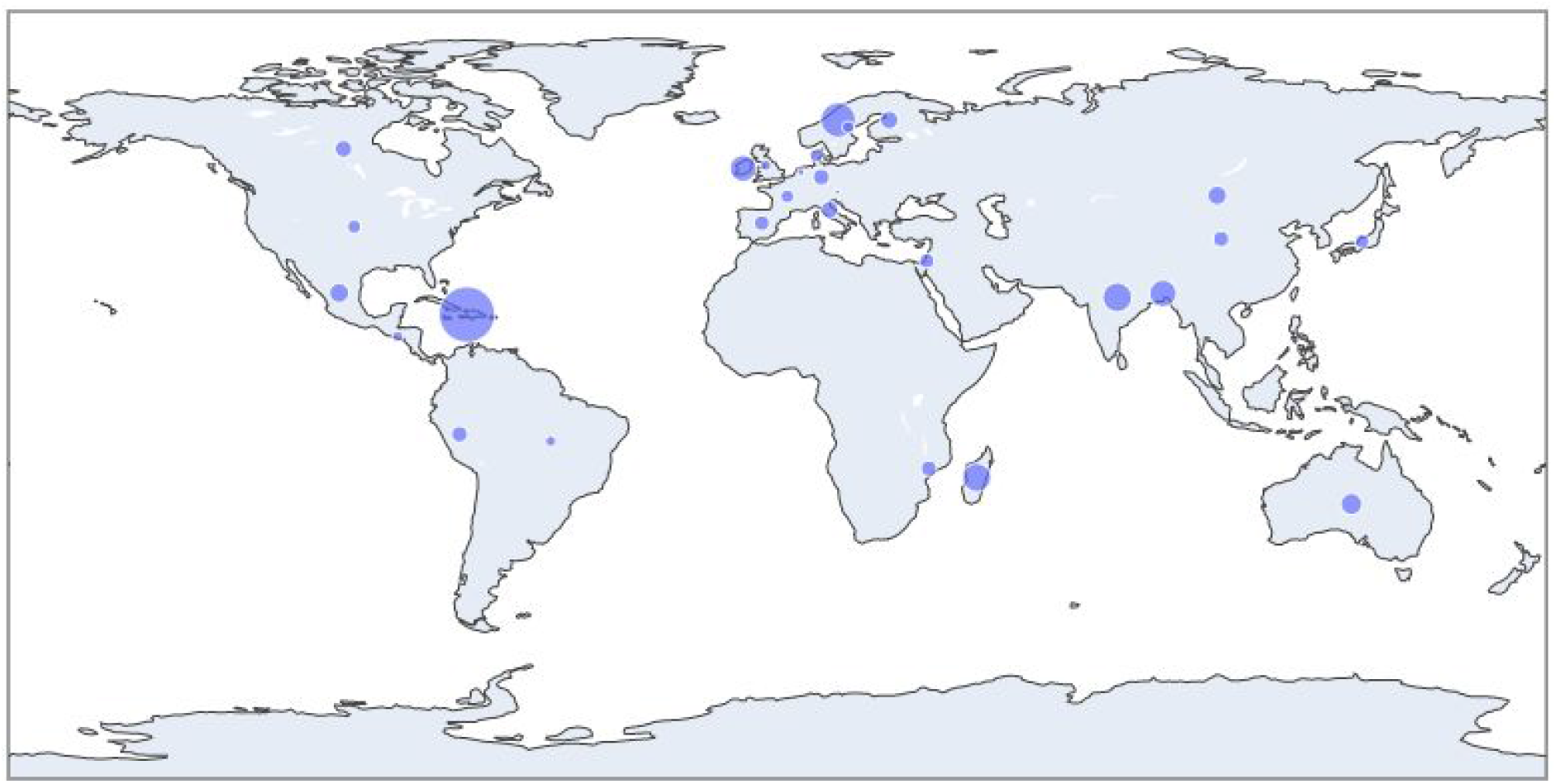
Relative prevalence of mcr-like genes (bubble area shows number of genes divided by number of genomes from each country) in countries with at least 50 genomes in the study.

The genus with the largest number of Mcr-like genes is *Sutterella* (667 genes), followed by *Parasutterella* (338 genes) and the alphaproteobacterial genus CAG-495 (258 genes) (Figure 2). The genus *Sutterella* is highly prevalent and, mainly represented by *S. wadsworthensis*, abundant in the healthy human gut^49^ and not considered pathogenic in general^50,51^. However, a role in autism in children was suggested^52^, and a higher prevalence in pre diabetics was observed^53^, while the role of *Sutterella* therein remains unknown. Due to isolations from several inflamed body parts, *S. wadsworthensis* might be considered an opportunistic pathogen^54^. While some AR occurred in *S. wadsworthensis*, so far, there was no report of Colistin resistance in the *Sutterella* genus. At least one study reported a *S. wadsworthensis* strain as susceptible^55^, indicating that colistin susceptibility of *Sutterella* strains from the human gut must be experimentally validated in the future. Similar to *Sutterella*, *Parasutterella* species, while sometimes linked to diseases, appear to be ordinary members of the human gut microbiota^56^, pathogenic only in rare cases. The unstudied CAG-495 genus was reported to be abundant in patients diagnosed with Vogt-Koyanagi-Harada disease^57^.

**Figure 2:**
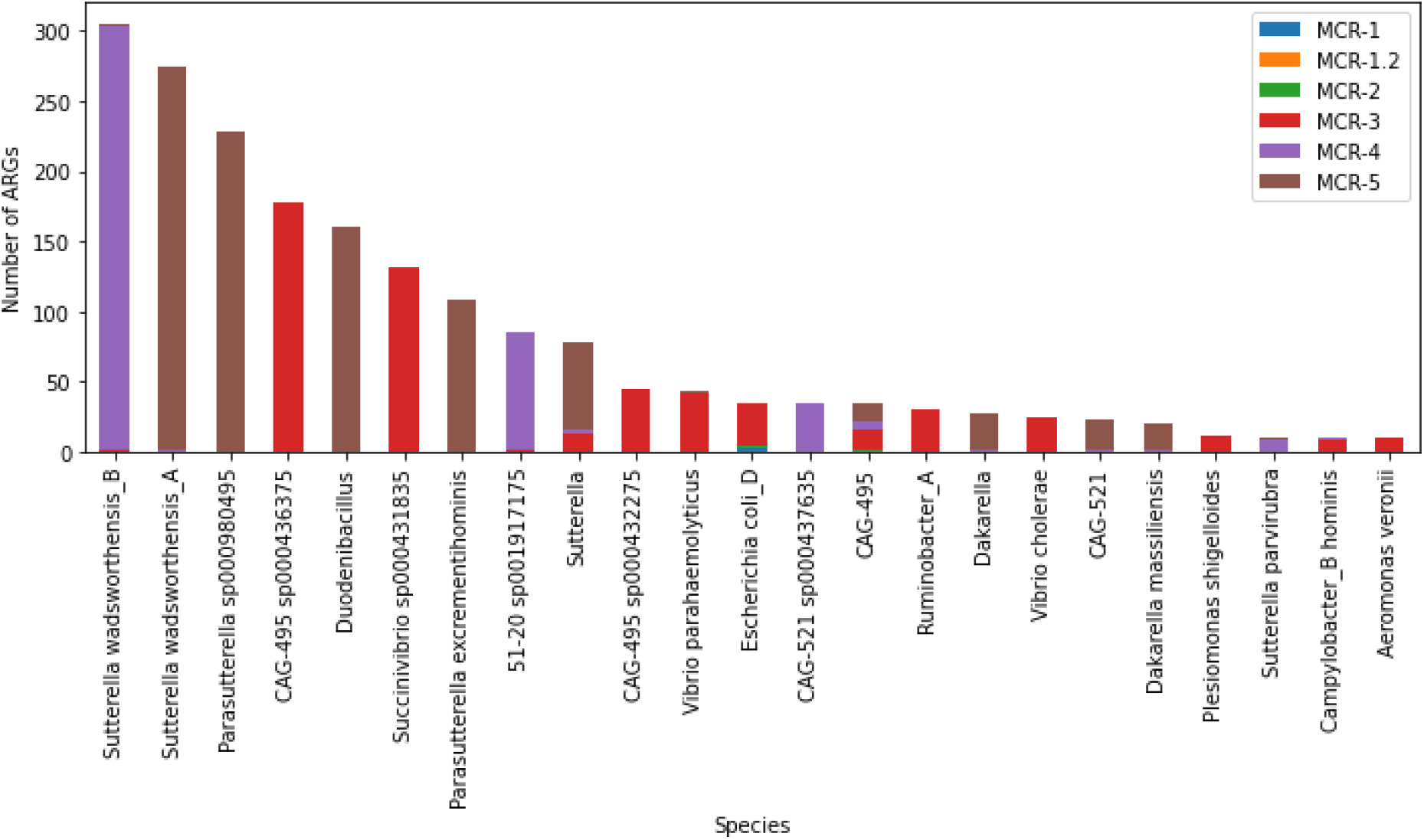
Prevalence of Mcr-like genes in different genera (with 10 or more Mcr-like genes).

The main potentially pathogenic species carrying Mcr-like genes belong to the genera *Vibrio*, *Escherichia* (only a few pathogenic strains, mainly a commensal gut bacterium), and *Campylobacter*. While the relative abundance of mcr-like genes in *Escherichia* is low (< 1%), that for Vibrio (>10%) and *Campylobacter* (>50%) is relatively high and warrants further investigation if the human gut might contribute to spreading colistin-resistant strains of these pathogens.

Many of the mcr-like gene sequences identified in this study are much larger than the average MCR (~530 amino acids long), with some reaching up to 831 amino acids. We found that there was an extra region encoding PAP2 and PAP2_like domains (an example can be found in Figure 3). In MCR-1 plasmids, PAP2 is encoded in a separated ORF downstream mcr sequence^58^.

**Figure 3:**
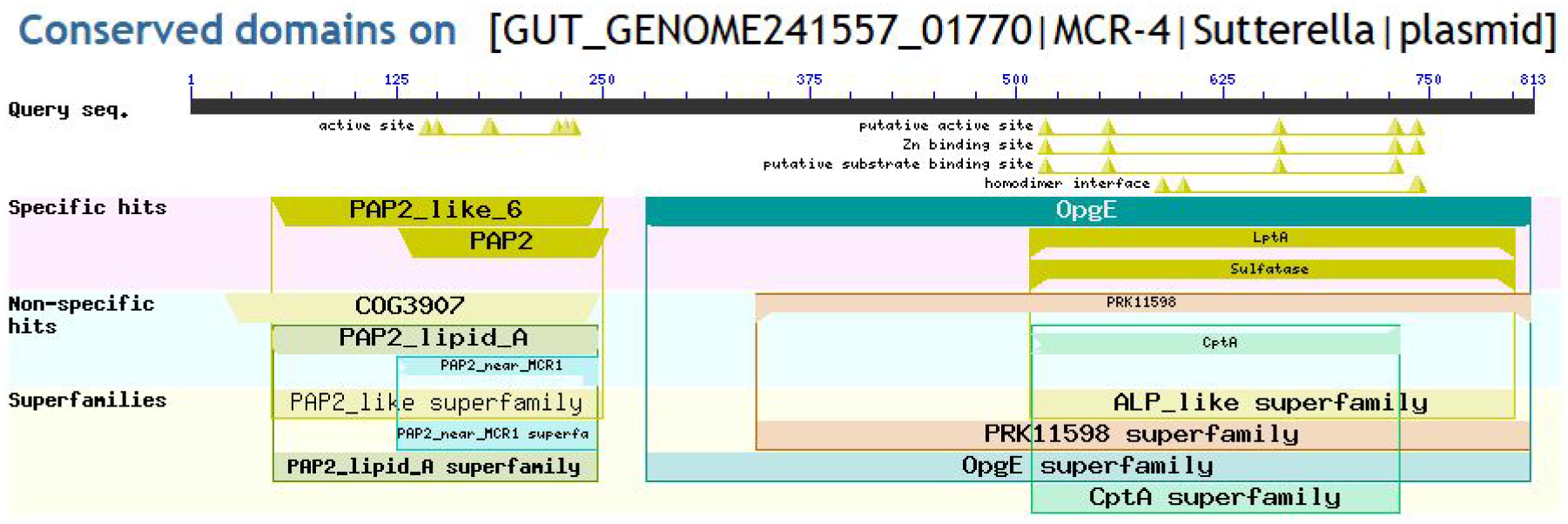
Conserved domains on one MCR-4 like sequence from the Sutterella genus. This analysis was done on CDD online from NCBI blast tool.

Regarding their genomic context, a total of 215 genes are present in contigs classified as plasmid by PlasFlow, while 1239 were in contigs classified as chromosomes, and 625 genes were in not classified contigs.

We also verified the existence of other ARGs detected by deepARG in genomes containing Mcr-like sequences. We identified a total of 22,746 ARGs (from 21 ARG classes and some unclassified) co-occurring with MCR-like sequences, being the most abundant classes the multidrug resistance (10,008 ARGs), beta-lactam (2,271 ARGs) and glycopeptide (2,261 ARGs) (Figure 4). However, several of those sequences on the multidrug class are efflux proteins, and as discussed in our previous study^9^, those are very hard to distinguish from other transporters that are not involved in antibiotic resistance.

**Figure 4:**
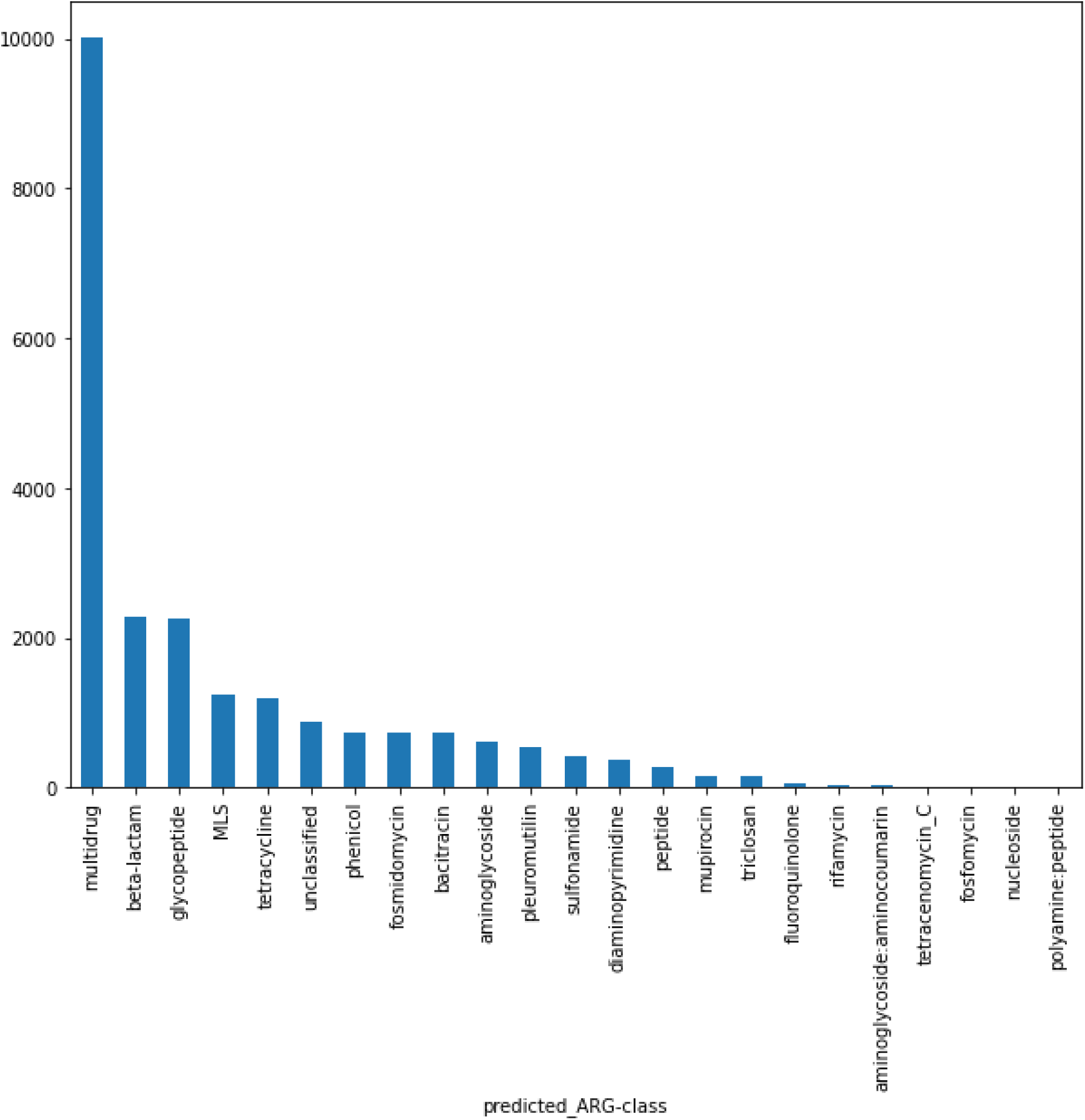
Number of ARGs per class co-occurring in the same genome with Mcr-like genes.

Colistin is a last resource antibiotic used against MDR bacteria with extended-spectrum beta-lactamases (ESBL), which makes the investigation of the presence of those ARGs in genomes containing MCR-like sequences so important. Regarding the beta-lactam group, we identified 1138 penA, a penicillin-binding protein 2 (PBP 2) associated with reduced susceptibility to oral cephalosporins^59^. Besides, we also identified 673 pbp1-A and 129 pbp1-B that are also penicillin-binding proteins.

Regarding the ESBL group, we identified 25 blaOXA (class D beta-lactamase capable of destroying 3rd generation cephalosporins)^60^, 3 blaCTX-M (a plasmid-encoded ESBL found in Enterobacteriaceae, likely acquired from the environmental bacteria *Kluyvera spp.* by HGT^61^), 2 blaTEM-153, 7 blaTLA-1 and 2 blaCFXA-6. The blaCTX-M enzymes have been found associated with insertion elements (ISEcp1) and transposable elements (for example, Tn402-like transposons). Many conjugative plasmids can transport these mobile elements, and consequently, these enzymes became the most prevalent ESBL ^62 63^.

The phylogenetic tree (Figure 5) shows the protein sequences classified as MCR-4 and MCR-5 in *Sutterella*, *Parasutterella*, and CAG-521 together in a clade with support value 1, and clinical MCR-5 sequences in an adjacent clade. The sequences used in the tree are grouped with clinical MCRs instead of the outgroup eptA, providing additional evidence for the annotation of those sequences.

**Figure 5:**
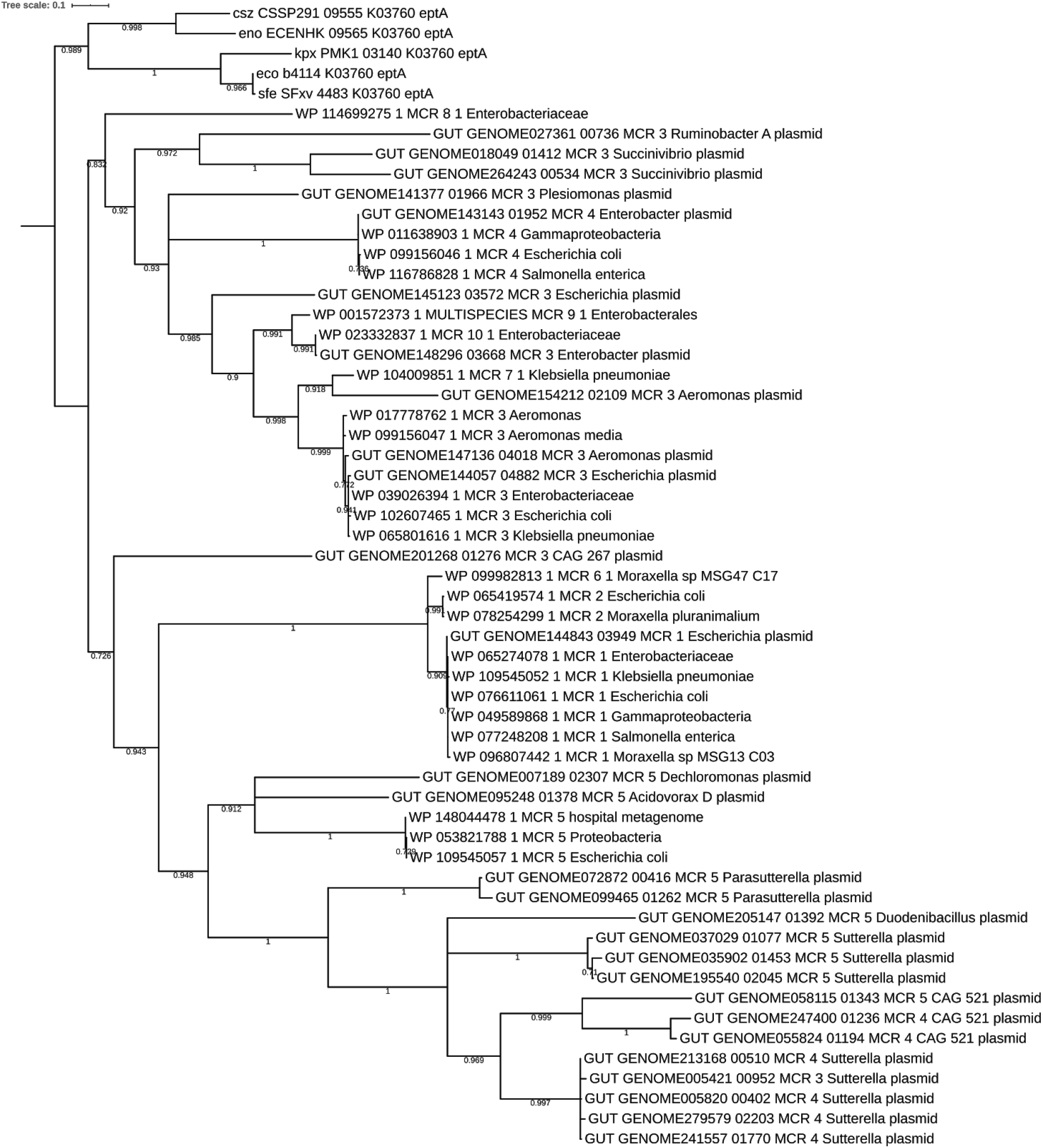
Phylogenetic tree of MCR-like sequences. We used only sequences present in contigs classified as plasmids and we clusterized similar sequences with CD-HIT on 97% similarity. The phylogenetic informative regions were selected by BMGE and the Maximum likelihood (ML) phylogenetic tree was calculated by PhyML 3.0 with the model selection performed by SMS (AIC method) and 100 bootstrap replicates to infer significance. We added clinical MCR sequences from NCBI to the analysis.

## Conclusions

This study uncovers the diversity of MCR-like genes in the human gut microbiome. We showed the cosmopolitan distribution of these genes in patients worldwide and the co-presence of other antibiotic resistance genes, including ESBLs. Also, we described mcr-like genes encoded in the same ORF with PAP2-like in bacteria from the genus Sutterella. Although these novel sequences increase our knowledge about the diversity and evolution of mcr-like genes, their activity has to be experimentally validated in the future.

## Data and Code availability

All the code used in this study is available at https://github.com/rcuadrat/human_microbiome_mcr and all the data is available at Zenodo (https://doi.org/10.5281/zenodo.4399676).

## Acknowledgment

This research has received funding from the European Union’s Horizon 2020 research and innovation programme under the Marie Skłodowska-Curie grant agreement No. 801522, by Science Foundation Ireland and co-funded by the European Regional Development Fund through the ADAPT Centre for Digital Content Technology grant number 13/RC/2106. We thank Maria Sorokina and also the colleagues in DIfE for the meaningful discussions during the elaboration of this text.

